# Assembly of continuous high-resolution draft genome sequence of *Hemicentrotus pulcherrimus* using long-read sequencing

**DOI:** 10.1101/2023.10.31.565072

**Authors:** Tetsushi Komoto, Kazuho Ikeo, Shunsuke Yaguchi, Takashi Yamamoto, Naoaki Sakamoto, Akinori Awazu

## Abstract

The update of the draft genome assembly of sea urchin, *Hemicentrotus pulcherrimus*, which is widely studied in East Asia as a model organism of early development, was performed using Oxford nanopore long-read sequencing. The updated assembly provided ∼600 Mb genome sequences divided into 2,163 contigs with N50 = 516 kb. BUSCO completeness score and transcriptome model mapping ratio (TMMR) of the present assembly were obtained as 96.5 % and 77.8 % respectively. These results were more continuous with higher resolution than those by the previous version of *H. pulcerrimus* draft genome, HpulGenome_v1, where the number of scaffolds = 16,251 with a total of ∼100 Mb gaps, N50 = 143 kb, BUSCO completeness score = 86.1 %, and TMMR = 40.61. The obtained genome contained 36,055 gene models that were consistent with those in other echinoderms. Additionally, two tandem repeat sequences of early histone gene locus containing 47 copies and 34 copies of all histone genes, and 185 of the homologous sequences of the interspecifically conserved region of the *Ars* insulator, ArsInsC, were obtained. These results provide further advance for genome-wide research of development, gene regulation, and intranuclear structural dynamics of multicellular organisms using *Hemicentrotus pulcherrimus*.

## INTRODUCTION

Sea urchins are model organisms commonly used for studying early development and morphogenesis due to its evolutionary position as diverged at the early period of deuterostome evolution. For example, *Hemicentrotus pulcherrimus* which is one of the typical sea urchin in East Asia was widely investigated to reveal developmental systems of multicellular organisms such as the intranuclear chromatin structural dynamics, establishment of left-right asymmetric body axis, and nervous system formation (Matsushita et al., 2017; Takemoto et al., 2016; Yaguchi & Katow, 2003). The gene regulatory network controlling endomesoderm specification in sea urchin embryos was also extensively studied using *Strongylocentrotus purpuratus* (E. H. Davidson et al., 2002; Oliveri & Davidson, 2004).

The detailed features of genome sequences of various organisms promote the research for a variety of universal and specific activities of the entire living systems. Therefore, as similar to various organisms, sea urchins draft genomes were also assembled; for example *Sp* in 2006 (Sodergren et al., 2006) (the latest version was updated in 2019), *H. pulcerrimus* in 2018 (Kinjo, Kiyomoto, Yamamoto, Ikeo, & Yaguchi, 2018), *Lytechinus variegatus* in 2020 (P. L. Davidson et al., 2020), *Temnopleurus reevesii* in 2022 (Kinjo et al., 2022), and *Paracentrotus lividus* in 2023 (Marletaz, Couloux, & Poulain, 2023).

Recent research by the Hi-C method and its derivatives (Erez Lieberman-Aiden et al., 2009), which has progressed rapidly in the last decade, has revealed that the expression of each gene is affected by various *cis*-regulatory elements and their higher-ordered structures such as enhancer-promoter loops, nucleosome exclusive non-looping insulator sequence (NENLIS) (Matsushima, Sakamoto, & Awazu, 2019), topologically associated domain (TAD), A/B compartment, etc. (E. Lieberman-Aiden et al., 2009). To carry out the analyses of the effects of such *cis*-regulatory structures at various scales ranging from several kb to Mb, a highly contiguous genomic sequence longer than several Mb of the target organism is required.

In contrast, it is difficult to construct such a long contiguous scaffold that contains a transcribed gene region and its *cis*-regulatory elements such as enhancers and insulators with conventional short-read-based genome assembly. For example, previously reported draft genome sequence of *H. pulcerrimus*, called HpulGenome_v1, have not contained the *Ars* insulator sequence locating at 2 kb upstream of arylsulfatase (*Hp-Ars)* gene (Akasaka et al., 1999; Takagi et al., 2012), whose core region (ArsInsC) is known as a typical NENLIS (Matsushima et al., 2019). Furthermore, *Hp-Ars* gene-containing scaffold in the HpulGenome_v1 did not include another important regulatory region, C15 enhancer (Iuchi et al., 1995; Sakamoto et al., 1997). Since repetitive sequences are dispersed throughout the genome, the presence of repeat sequences leads to difficulty in the short-read-based genome assembly. In addition, it was impossible to reconstruct long repeat sequences far exceeding several hundred bp, which frequently appear in multicellular organism genomes, and therefore, many gaps had to exist in the scaffold. Thus, although it has been suggested that the *H. pulcerrimus* genome contains long repetitive sequences consisting of tens to hundreds of tandem sequences of genes encoding histones H1, H2A, H2B, H3, and H4, HpulGenome_v1 does not contain such long repeat regions.

However, the above problems are being overcome by technological advances in long-read sequencing, which yields sequences of tens to hundreds of kb per read. At present, reads obtained by long-read sequencing appear to contain errors 100 times more frequently than those obtained by short-read sequencing. However, the hybrid assembly method, which assembles short reads from the same organism while correcting errors in long-reads, may enable highly accurate genome assembly with long continuity by taking advantage of both.

Therefore, in this study, the *H. pulcerrimus* draft genome sequence was updated by hybrid assembly using both long-reads obtained using the Oxford Nanopore sequencer and short-reads used in conventional *H. pulcerrimus* draft genome assembly. In this paper, we compared the updated draft genome sequence with HpulGenome_v1, and provided the annotation of genes by comparison with related species. We would also show that the updated draft genome contains ArsInsC sequence at the upstream of *Hp-Ars* gene and its homologous sequences in the updated genome. Furthermore, we identified two distinct long tandem repeats of early histone genes. Such improvement of the draft genome sequence will drive further progress of studies for physiological and structural-dynamical features of gene regulatory behaviors in *H. pulcherrimus*.

## MATERIALS AND METHODS

### Sample Collection and DNA Extraction

Adult males of *H. pulcherrimus* were collected from the sea around Etajima City, Hiroshima Prefecture, Japan under the permission of Hiroshima Fishery Cooperative, and extracted genomic DNA (gDNA) from their sperm. The *H. pulcherrimus* genomic DNA was extracted by using Blood & Cell Culture DNA Midi Kit (QIAGEN), and short genomic DNA segments were removed by Short Read Elimination XL Kit (Circulomics Inc.). To obtain extremely long-reads, we followed the protocol described by Logsdon (2020).

### Sequencing Library Preparation

A Nanopore sequencing libraries were prepared from these gDNA using Rapid Sequencing Kit (SQK-RAD004; Oxford Nanopore Technologies [ONT]). Nanopore sequencing were performed on an ONT MinION sequencer using R9.4.1 flow cells and FAST5 files were basecalled to FASTQ files using ONT MinIT (MNT-001). In the following, each of the generated fastq formatted reads was named ONT-read.

### Preprocessing of Illumina short-reads and ONT-read Data

The preprocessing (adapter trimmed and removed low-quality reads) of Illumina short-reads from gDNA of *H. pulcherrimus* (DRR107784, DRR107785 and DRR107786) and transcriptome (DRR107783) were performed using fastp v0.22.0 (9). Error correction of ONT-read data was performed using Ratatosk v0.7.0 (Holley et al., 2021) with preprocessed Illumina short-reads (160.47 Gb in total).

### Genome Assembly and Polishing

De novo genome assembly for *H. pulcherrimus* were performed using over 5 kb error-corrected ONT-reads by Raven v1.8.1 (Vaser & Šikić, 2021), Flye v2.8.3-b1695 (Kolmogorov, Yuan, Lin, & Pevzner, 2019) and Wtdbg2 v2.5 (Ruan & Li, 2020). These draft genome sequences were corrected with each other and a single consensus sequence was generated using MAECI (Lang, 2022). This consensus sequence was polished with Illumina short-reads using MAECI.

### Assessment of Draft Genome Sequences

To evaluate the completeness of genome assembly, searching for highly conserved genes in metazoan using BUSCO v5.3 (Manni, Berkeley, Seppey, & Zdobnov, 2021). As an additional analysis, predicted transcriptome models in HpBase were mapped to each draft genome sequence using hisat2 v2.2.1 (Kim, Paggi, Park, Bennett, & Salzberg, 2019).

### Gene Prediction and Gene Annotation

The preprocessed Illumina short-reads from transcriptome were mapped to draft genome sequences using hisat2 v2.2.1 (Kim et al., 2019), and generate BAM formatted files using SAMtools v1.17 (Li et al., 2009). Gene prediction was performed using these BAM files by BRAKER2 v2.1.4 (Brůna, Hoff, Lomsadze, Stanke, & Borodovsky, 2021). Gene annotation (orthology search) of gene models predicted by BRAKER2 was performed using BLASTP v2.6.0 (Altschul, Gish, Miller, Myers, & Lipman, 1990) with protein models in HpBase and Echinobase (*Strongylocentrotus purpuratus* v5.0 and *Lytechinus variegatus* v3.0).

## RESULTS

### Genome Assembly and Completeness of Draft Genome Sequence

The sequencing by ONT MinION sequencer generated 17.29 M reads with 51.54 Gb sequenced data. By the error correction and filtering out ONT-reads, 2.76 M reads with 35.26 Gb were generated for genome assembly. The draft genome sequence was updated by assembling and polishing these reads by Raven, Flye, Wtdbg2 and MAECI (Table 1. Table S1). Here, the contig (contig ID: Utg196084) that was most homologous to the mitochondrial genome of *Hemicentrotus pulcherrimus* [NC_023771.1] by BLASTN was removed. The updated draft genome sequence consists of 2,163 contigs with a total length ∼ 626.4 Mb and N50 length ∼ 515.7 kb (Fig. 1A, Table S1), that contained smaller number of contigs with more continuously assembled than HpulGenome_v1; HpulGenome_v1 consists of ∼ 16,000 scaffolds (contigs) with total length ∼ 600 Mb containing totally ∼100 Mb gaps and N50 = 142.6 kb (Fig. 1B).

**TABLE 1.** Summary of genome assembly, BUSCO completeness and gene annotation in HpulGenome_v1 and our draft genome sequence. * were calculated from BBtools (Bushnell, Rood, & Singer, 2017) (stats.sh).

|  | *Updated draft genome* | *HpulGenome_v1* |
| --- | --- | --- |
| Assembly size* | 626.4 Mb | 568.9 Mb |
| No. contigs* | 2,163 | 86,128 |
| N50 contig length* | 515.7 kb | 9.641 kb |
| No. scaffolds* | 2,163 | 16,251 |
| N50 scaffold length* | 515.7 kb | 142.6 kb |
| N (%)* | 0 | 16.41 |
| GC-content (%)* | 37.12 | 36.72 |
| BUSCO completeness (%) | Complete : 96.5 | Complete : 86.1 |
| (metazoa_odb10: 954 genes) | Duplicated : 6.6 | Duplicated : 1.3 |
|  | Fragmented : 2.1 | Fragmented : 9.9 |
|  | Missing : 1.4 | Missing : 4.0 |
| Mapping ratio | 75.65 (aligned exactly 1 time) | 54.64 (aligned exactly 1 time) |
| of transcriptome models (%) | 2.13 (aligned >1 times) | 0.71 (aligned >1 times) |
| (20,564 sequences) |  |  |

**Fig 1.**
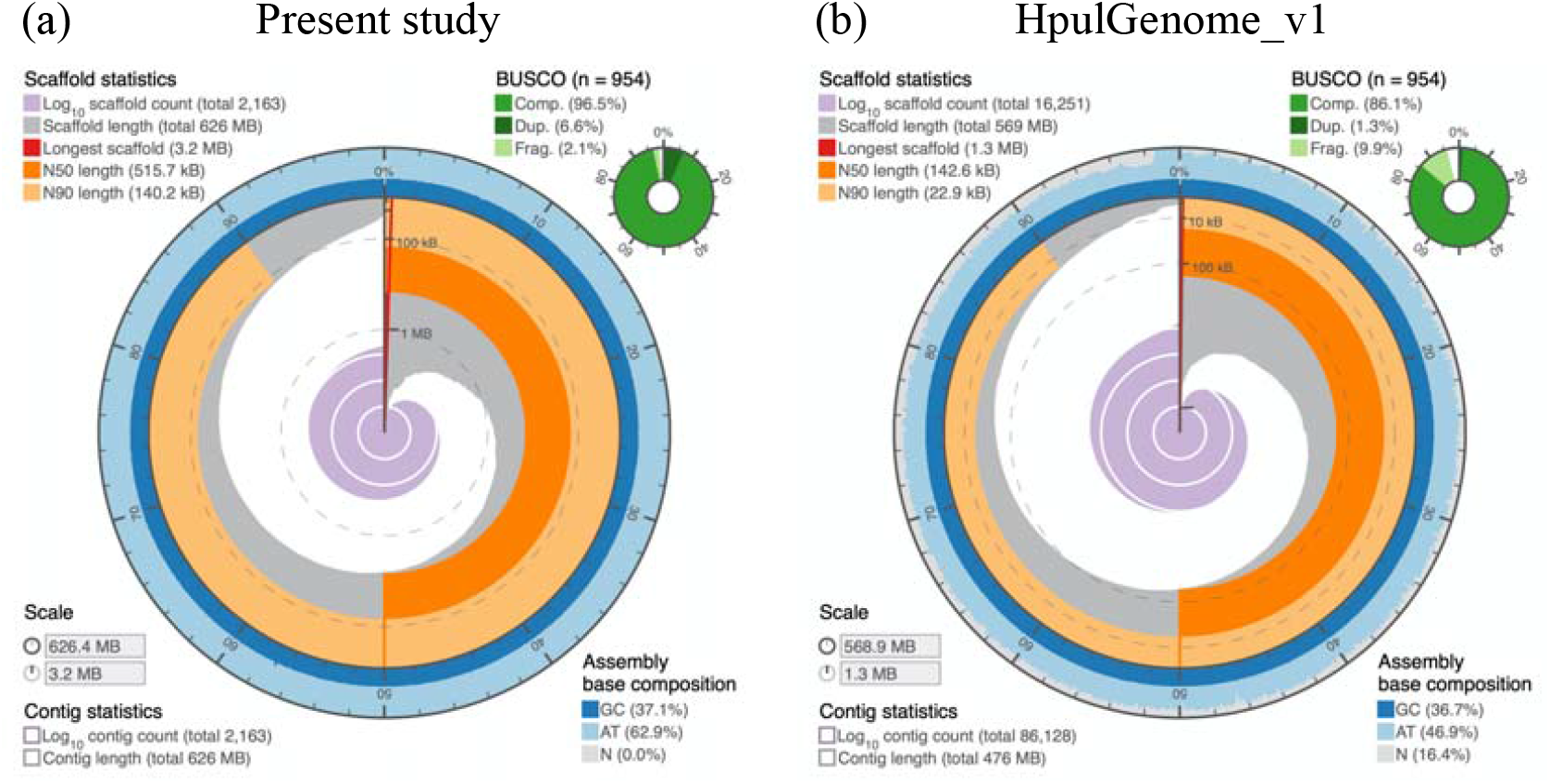
Assembly metrics of the obtained draft genome sequence obtained in the present study (a), and that of HpulGenome_v1 (b). Assembly metrics were visualized using assembly-stats. (https://github.com/rjchallis/assembly-stats)

Genome completeness analysis of the updated draft genome by BUSCO using the metazoa_odb10 showed larger value as 96.5 % (89.9 % single-copy and 6.6 duplicated) than that of HpulGenome_v1, 86.1 % (84.8 % single-copy and 1.3 duplicated). Additionally, the mapping ratio of transcriptome models of the present draft genome was also obtained larger value as 77.78 % than that of HpulGenome_v1, 55.35 %.

### Gene Prediction Based on Genome Assembly and Gene Annotation with Related Species

The analysis by BRAKER2 and transcriptome data (DRR107783) predicted 46,914 genes exist in the updated draft genome sequence. By the elimination of the predicted genes that was too short (less than 50 amino acids), contained multiple stop codons, or was encoded on the mitochondrial contig, 46,826 gene models were obtained in the updated draft genome (Table S2). The inference from the orthology relationships between the present gene models and gene models in other sea urchins through the reciprocal BLASTP searches showed that 36,055 gene models (# of reciprocal best hit pair: 20,434, and # of not reciprocal but best hit: 15,621) were homologous (best hit with e-value ≤ 1e-10) to publicly known gene models of *H. pulcerrimus, S. purpuratus* or *L. variegatus*

(Table 2, Table S2).

**TABLE 2.** Summary of BLASTP searches with protein models of HpBase and other sea urchin. Proteins fasta file of *Paracentrotus lividus* was created using gffread v0.12.7 (Pertea & Pertea, 2020). The genome and annotation in GTF format of *P. lividus* were downloaded from Zenodo (https://zenodo.org/record/7459274).

| Subject | Reciprocal | Not reciprocal | No hit |
| --- | --- | --- | --- |

TABLE 2. Summary of BLASTP searches with protein models of HpBase and other sea urchin.
|  | best hit pair | but best hit |  |
| --- | --- | --- | --- |
| ***H. pulcherrimus***<br>**(HpBase,**<br>**HpulGenome_v1_prot.fa)** | 15,652<br>(33.43%) | 16,847<br>(35.98 %) | 14,327<br>(30.60 %) |
| ***S. purpuratus***<br>**(Echinobase,**<br>**sp5_0_GCF_proteins.fa)** | 16,153<br>(34.50 %) | 17,190<br>(36.71 %) | 13,483<br>(28.79 %) |
| ***L. variegatus***<br>**(Echinobase,**<br>**Lvar3_0_GCF_proteins.fa)** | 14,802<br>(31.61 %) | 16,613<br>(35.48 %) | 15,411<br>(32.91 %) |
| ***P. lividus*\*** | 14,958<br>(31.94 %) | 18,388<br>(39.27 %) | 13,480<br>(28.70 %) |
| ***H. pulcherrimus, S. purpuratus,***<br>***or L. variegatus*** | 20,434<br>(43.64 %) | 15,621<br>(33.36 %) | 10,771<br>(23.00 %) |

### Detection of Early Histone Gene Loci with Long Tandem Repeats

*H. pulcherrimus* genome has been suggested to contain two non-allelic early histone gene loci that are the long repetitive sequences consisting of tens to hundreds of tandem sequences of genes encoding histones (Table S3) (Matsushita et al., 2017). However, such long tandem repeat sequence could not be detected by recent short-read-based genome assemblies; In HpulGenome_v1, BLASTN searches did not find any genomic regions homologous even to single repeat sequences of early histone loci.

In contrast, two contigs containing the regions of long tandem repeats of histone genes were found on the updated draft genome sequence; where one contig (contig ID: Utg198178, with 479,333 bp) involved 47 tandem repeats, and the other contig (contig ID: Utg200276, with 127,611 bp) involved 34 tandem repeats (Table S3).

### Regulatory sequences of arylsulfatase gene

We have investigated the regulatory mechanism of the transcription of the *arylsulfatase* (*Hp-Ars*) gene. The promoter (−252 to +38) is the minimum region required for temporal expression (Iuchi, Morokuma, Akasaka, & Shimada, 1995; Morokuma, Akasaka, Mitsunaga-Nakatsubo, & Shimada, 1997). The C15 enhancer required for large enhancement of the expression is present in the first intron (Iuchi et al., 1995; Sakamoto et al., 1997). Polypyrimidine sequence (−2,201 to -1,680) that can form an intramolecular triplex structure (Sakamoto, Akasaka, Yamamoto, & Shimada, 1996; Yamamoto, Akasaka, Irie, & Shimada, 1994) and the *Ars* insulator (−2,686 to -2,109) (Akasaka et al., 1999) are present in the upstream region.

In HpulGenome_v1, scaffold989 contains the *Hp-Ars* gene. BLASTN search revealed that this scaffold includes all exons, the promoter, and upstream sequences to the polypyrimidine region. However, it does not include the C15 enhancer in the 1st intron and the *Ars* insulator in the upstream flanking region, which play important roles in the transcriptional regulation.

On the other hand, in the updated draft genome, *Hp-Ars* gene-containing contig (contig ID: Utg200732) contains all exons and regulatory sequences. Furthermore, *Hp-Ars* gene has been reported to include various types of repetitive sequences such as direct repeats and inverted repeat (Akasaka et al., 1994). All of these repetitive sequences could be found in the contig Utg200732. Interestingly, when we searched homologous sequences of the inverted repeat (−475 to -217) with 90-120 bp of the unit length, >90% identity, and >90% query coverage among inverted repeat obtained from IRF v3.08 (Warburton, Giordano, Cheung, Gelfand, & Benson, 2004), 9 homologous inverted repeat sequences were found in the updated draft genome (Table S4). Although 7 out of these homologous inverted repeats were present in intergenic regions, there was no tendency in their position. Furthermore, 8 homologous sequences of the direct repeat DIR1 (−3,440 to -3,109) with 2 repeat unit, >90% identity, and >95% query coverage were found, and 205 homologous sequences of another direct repeat DIR2 (−3,096 to -2,592) with 2-5 repeat unit, >90% identity and >95% query coverage were found in the updated draft genome (Table S5). These DIR1 and DIR2 homologs tend to exist in the upstream region of the nearest genes (Fig. 2).

**Fig 2.**
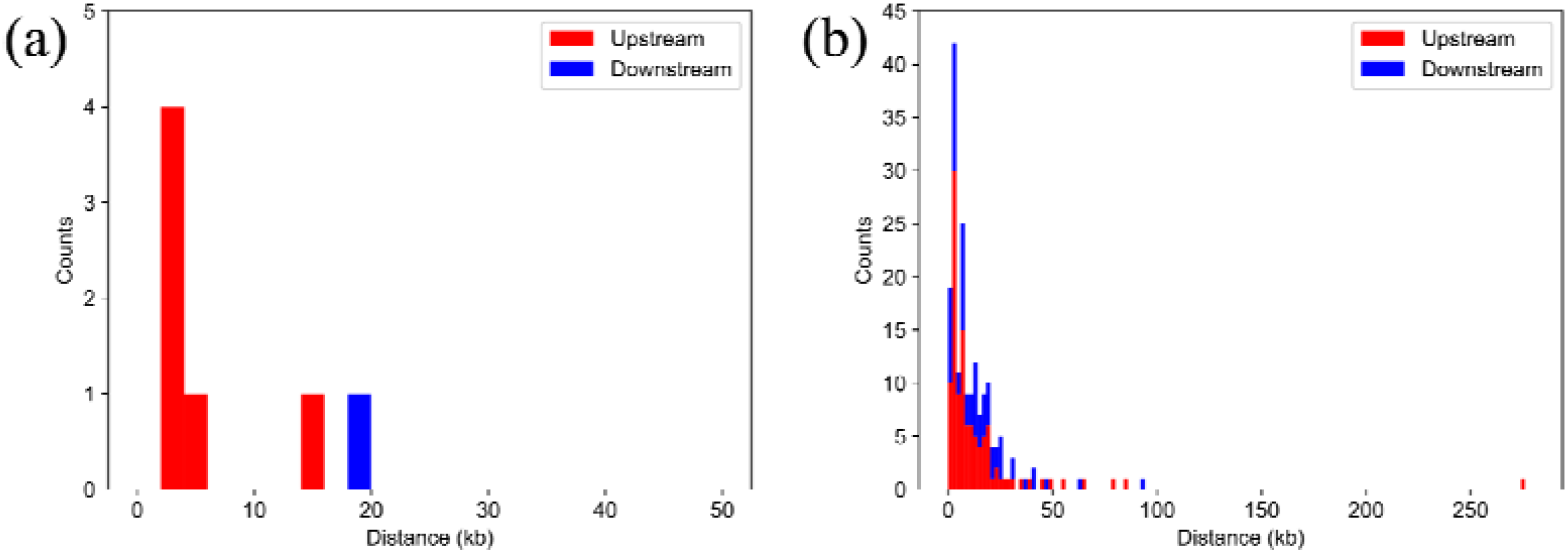
Stacked histogram of genomic distances between DIR1 homologs and their nearest gene (a), and that between DIR2 homologs and their nearest gene (b). Height of red and blue bars indicates counts of DIR homologs at the upstream and downstream region of the nearest gene, respectively. Distances between DIR homolog and its nearest gene was defined as the genomic distance between the centers of them.

### Homologous genome region to *Ars* insulator

The *Hp-Ars* gene has been known to involve the *Ars* insulator, which includes interspecifically conserved ArsInsC sequence (Akasaka et al., 1999; Takagi et al., 2012) (Table S4) that is a typical example of NENLIS (Matsushima et al., 2019). In HpulGenome_v1, *Hp-Ars* gene was present, but ArsInsC sequence was not present. In contrast, the updated draft genome contained ArsInsC from 46,417 to 46,236 (minus-strand) in contig ID: Utg200732 where is the 2 kb upstream of the *Hp-Ars* gene region. As a result of BLASTN analysis, we found 185 sequences that have more than 90% identity and the same length as ArsInsC (Table S6). As observed in the *Ars* insulator, in 50 bp region surrounding the ArsInsC homologous sequences tended to have high GC contents (50-80% GC) (Fig. 3, Fig. S1), and 121 homologous sequences (65.4%) contained G(C)-stretch of more than 8 bp in these GC-rich regions (Fig. S1). Thus, the sequence properties of the *Ars* insulator are well conserved in these homologous sequences, suggesting that these homologous sequences may function as insulators in the sea urchin genome.

**Fig 3.**
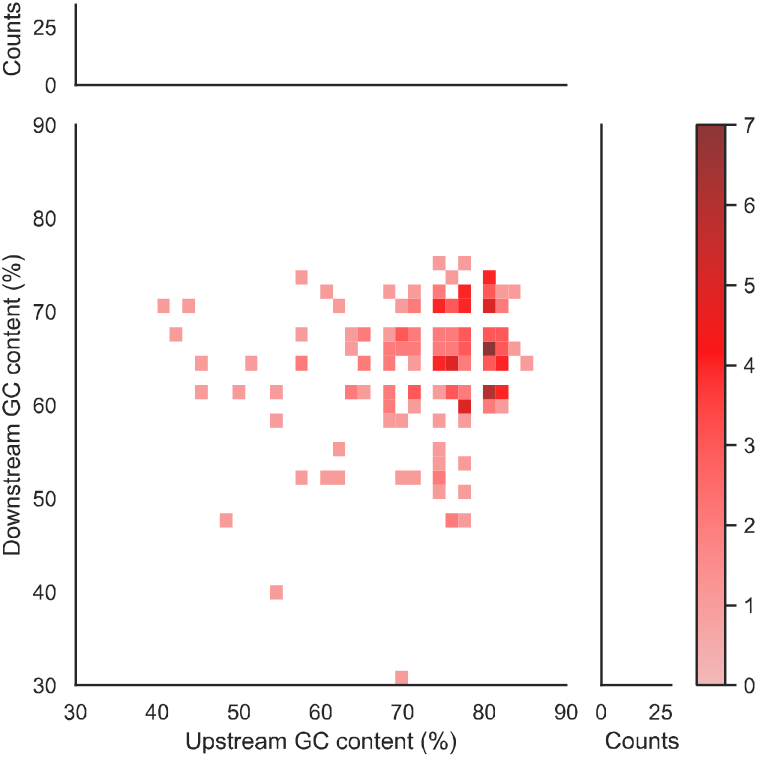
Heat map of the number of ArsInsC homologs with each GC content in the upstream 50bp and downstream 50bp regions. The histograms represent the projection onto the upstream GC content axis and the downstream GC content axis, respectively.

### Short tandem repeat (STR)

It has been reported that short tandem repeats (STRs) are enriched in gene promoters and variable repeat number is associated with the gene expression (Sawaya et al., 2013; Vinces, Legendre, Caldara, Hagihara, & Verstrepen, 2009). Furthermore, approximately 90% of transcription factors preferentially bind to STRs (Horton et al., 2023). Therefore, we also analyzed the presence of STRs in the updated draft genome using TRF v4.09 (Benson, 1999). According to the definition of microsatellite in Sawaya et al. (2013), uninterrupted consecutively repeated units of one to four nucleotides that are at least 12 bp in length were searched. After searching STRs that obtained from TRF results and matched this definition, a total of 111,553 mononucleotide repeats, 66,665 dinucleotide repeats, 61,983 trinucleotide repeats, and 62,448 tetranucleotide repeats were found in the updated draft genome (Table S7). However, the enrichment of STRs in the proximity of genes or the translation start sites was not observed.

## DISCUSSION

Long-read sequencing and hybrid assembly using long and short reads were performed to update *H. pulcherrimus* draft genome with more continuous and higher accuracy than recently reported one, HpulGenome_v1. This study resulted in the draft genome of *H. pulcherrimus* with higher values of the metrics by BUSCO and higher mapping rate of the transcriptome model than HpulGenome_v1.

This draft genome involved 46,826 gene models that is larger than HpulGenome_v1 containing 24,860 model genes. Here, 36,055 of these gene models were annotated based on the analysis with the homologous gene models of related sea urchins.

The present draft genome contained two long repeat sequences including the tandemly connected early histone genes. This result may drive the progress of the genomic and epigenomic analyses to reveal the developmental stage-dependent changes in intra-nuclear location and interactions of the early histone loci (Matsushita et al., 2017). Additionally, this draft genome also contained ArsInsC sequence and more than 100 of ArsInsC homologous sequences. In vertebrates, the CTCF-binding sequence is known as a typical insulator sequence responsible for gene expression control. Recently, however, it has been suggested that sea urchin CTCF functions during mitosis rather than interphase (Watanabe et al., 2023). Therefore, ArsInsC and its homologous sequences may play an important role in the control mechanism of sea urchin interphase gene expression, and the present draft genome may promote research to clarify the mechanism.

The updated draft genome involved more than 10,000 non-annotated genes. These genes were expected to include non-coding RNA-derived or *H. pulcherrimu* specific genes. The annotation of these gene models should be progressed by the comparison among genes in various organisms and the function analysis by experiments in the future. Additionally, the large number of duplicated genes were found by the analysis using BUSCO genes in the present draft genome than HpulGenome_v1. The reason for this fact was not clear whether each duplication is true in *H. pulcherrimu* or not. The further validation should be a future issue.

In this study, HpulGenome_v1 was updated using Oxford nanopore long-read sequencing, resulting in a smaller number of contigs with more continuous assembly. Since each contig became likely to contain a complete set of *cis*-regulatory sequences, we believe that this updated draft genome improves the applicability of *H. pulcherrimus* genome information to various genome-wide analysis.

## DATA AVAILABILITY

The sequence data of the updated genome assembly were featured on HpBase (https://cell-innovation.nig.ac.jp/Hpul/) with the name HpulGenome_kure_v1. The raw sequencing data of whole genome sequencing was deposited in the DDBJ Sequence Read Archive (DRA) with Accessions DRA017089, and was also submitted to DDBJ/EMBL/GenBank databases (BioProject Accession PRJDB16611)..

## Acknowledgements

This work was supported by a Grant-in-Aid for Scientific Research (KAKENHI) [Grant Number 21K06124] from the Japan Society for the Promotion of Science. Computations were partially performed on the NIG supercomputer at ROIS National Institute of Genetics.

