## Supplementary material for "Assembly of continuous high-resolution draft genome sequence of *Hemicentrotus pulcherrimus* using long-read sequencing": HP_ReAssembly_sup_FIN.docx

Supplementary information

Table S1: Final updated draft genome sequence (FASTA format) (a), and features of respectively obtained assembly results by Raven, Flye, and Wtdbg2 (b).

(a) “HpulGenome_kure_v1_contig.fa” in “https://cell-innovation.nig.ac.jp/cgi-bin/Hpul_public/Hpul_annot_download.cgi”

(b)

|  | Raven | Flye | Wtdbg2 |
| --- | --- | --- | --- |
| Assembly size* | 619.2 Mb | 985.5 Mb | 851.0 Mb |
| No. contigs* | 2,164 | 22,260 | 16,967 |
| N50 contig length* | 508.4kb | 144.8 kb | 240.3 kb |
| No. scaffolds* | 2,164 | 22,175 | 16,967 |
| N50 scaffold length* | 508.4 kb | 146.1 kb | 240.3 kb |
| N (%)* | 0 | 0 | 0 |
| GC-content (%)* | 36.91 | 37.36 | 36.94 |
| BUSCO completeness (%)  (metazoan_odb10:  954 genes) | Complete : 96.1  Duplicated : 6.5  Fragmented : 1.8  Missing : 1.8 | Complete : 96.6  Duplicated : 37.5  Fragmented : 2.6  Missing : 0.8 | Complete : 90.3  Duplicated : 10.4  Fragmented : 3.8  Missing : 5.9 |
| Mapping ratio of  transcriptome models (%)  (20,564 sequences) | 71.93 (aligned exactly 1 time)  3.12 (aligned >1 times) | 69.48 (aligned exactly 1 time)  8.00 (aligned >1 times) | 59.26 (aligned exactly 1 time)  1.85 (aligned >1 times) |

Table S2: Gene models and their features obtained by present assembled draft genome.

Nucleotide and amino acid sequences of each gene model (FASTA format) (a) and (b), gene transfer format (GTF) description of each gene position and feature (c), and gene annotation (d).

(a) “HpulGenome_kure_v1_nucl.fa” in “https://cell-innovation.nig.ac.jp/cgi-bin/Hpul_public/Hpul_annot_download.cgi”

(b) “HpulGenome_kure_v1_prot.fa” in “https://cell-innovation.nig.ac.jp/cgi-bin/Hpul_public/Hpul_annot_download.cgi”

(c) “https…/HpulGenome_kure_v1.gtf” in “https://cell-innovation.nig.ac.jp/cgi-bin/Hpul_public/Hpul_annot_download.cgi”

(d) “HpulGenome_kure_v1_annot.xlsx” in “https://cell-innovation.nig.ac.jp/cgi-bin/Hpul_public/Hpul_annot_download.cgi”

Table S3: Sequence of early histone locus (a), and locations of long repeated histone genes in HpulGenome_v1 and updated draft genome (b-d). Raw results obtained by BLASTN search for HpulGenome_v1(b) and updated draft genome (c), and locations of units of repeated histone genes (d).

(a) “HpEarlyHistone.fa”

(b) “HpulGenome_v1_BLASTN_withEarlyHistoneLoci.csv”

(c) “HpulGenome_kure_v1_BLASTN_withEarlyHistoneLoci.csv”

(d) “HpulGenome_kure_v1_EarlyHistoneLoci_47_34copies.csv”

Table S4: Sequence of *Ars*-INV (a) and subsequence used for BLASTN search (FASTA format) (b), and locations of homologous sequences to *Ars*-INV in updated draft genome (c). Raw results obtained by BLASTN search (d).

(a) “HpArs-INV.fa”

(b) “HpArs-INV_left.fa”

(c) “HpulGenome_kure_v1_HpArs-INV_homologue.csv”

(d) “HpulGenome_kure_v1_BLASTN_withHpArs-INV.csv”

Table S5: Subsequences of DIR1 (a) and DIR2 (b) (FASTA format), and location of homologous sequences to DIR1 (c) and DIR2 (d) in updated draft genome. Raw results obtained by BLASTN search of DIR1 (e) and DIR2 (f).

(a) “HpArs-DIR1.fa”
(b) “HpArs-DIR2.fa”
(c) “HpulGenome_kure_v1_HpArs-DIR1_homologue.csv”

(d) “HpulGenome_kure_v1_HpArs-DIR2_homologue.csv”

(e) “HpulGenome_kure_v1_BLASTN_withHpArs-DIR1.csv”

(f) “HpulGenome_kure_v1_BLASTN_withHpArs-DIR2.csv”

Table S6: Sequence of ArsInsC (a), locations of homologous sequences to ArsInsC in updated draft genome (b). Raw result obtained by BLASTN (c).

(a) “HpArsInsC.fa”

(b) “HpulGenome_kure_v1_ArsInsC_homologue.csv”

(c) “HpulGenome_kure_v1_BLASTN_withArsInsC.csv”

Table S7: Locations of STRs in updated draft genome.

“HpulGenome_kure_v1_ShortTandemRepeats.csv”

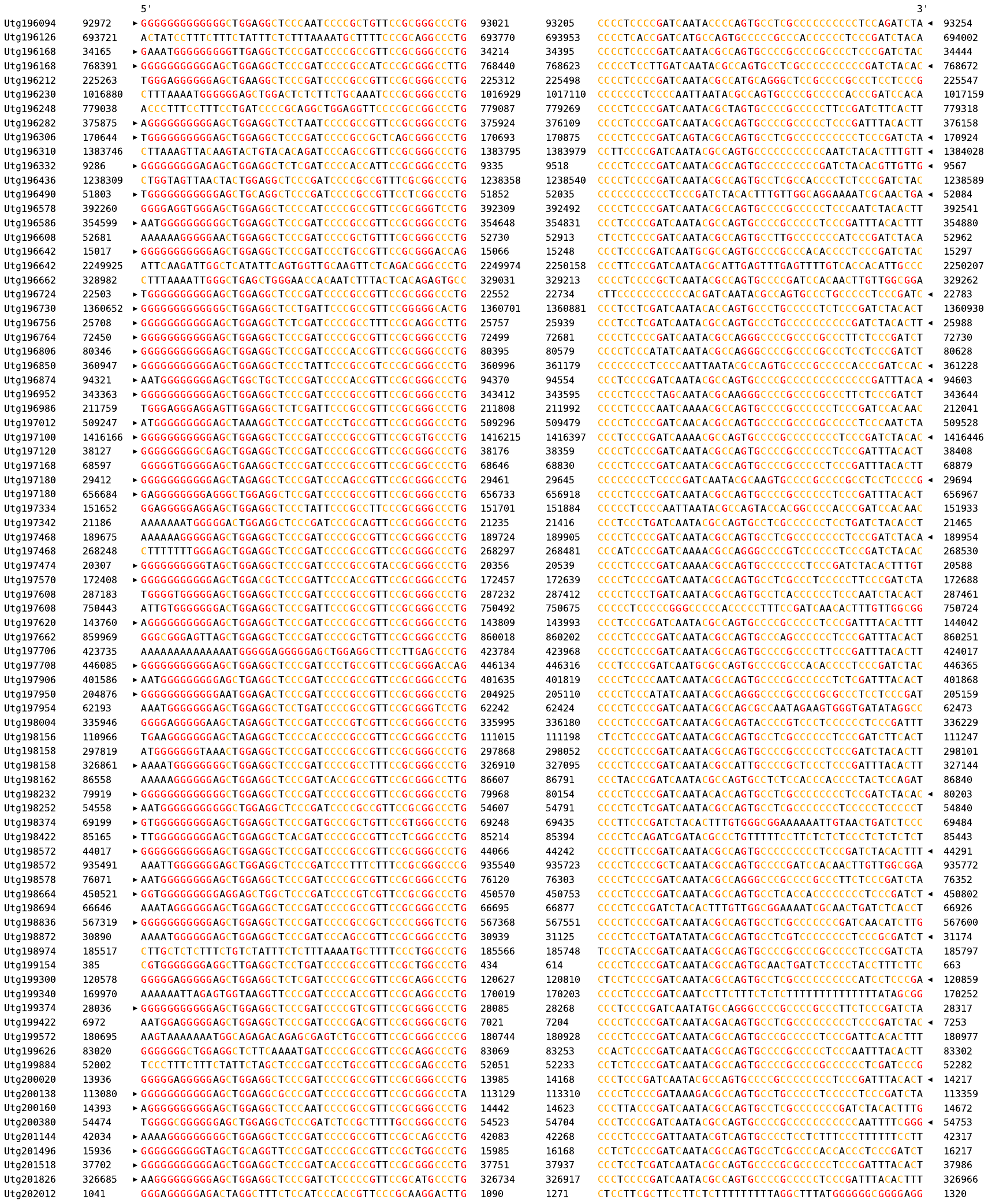

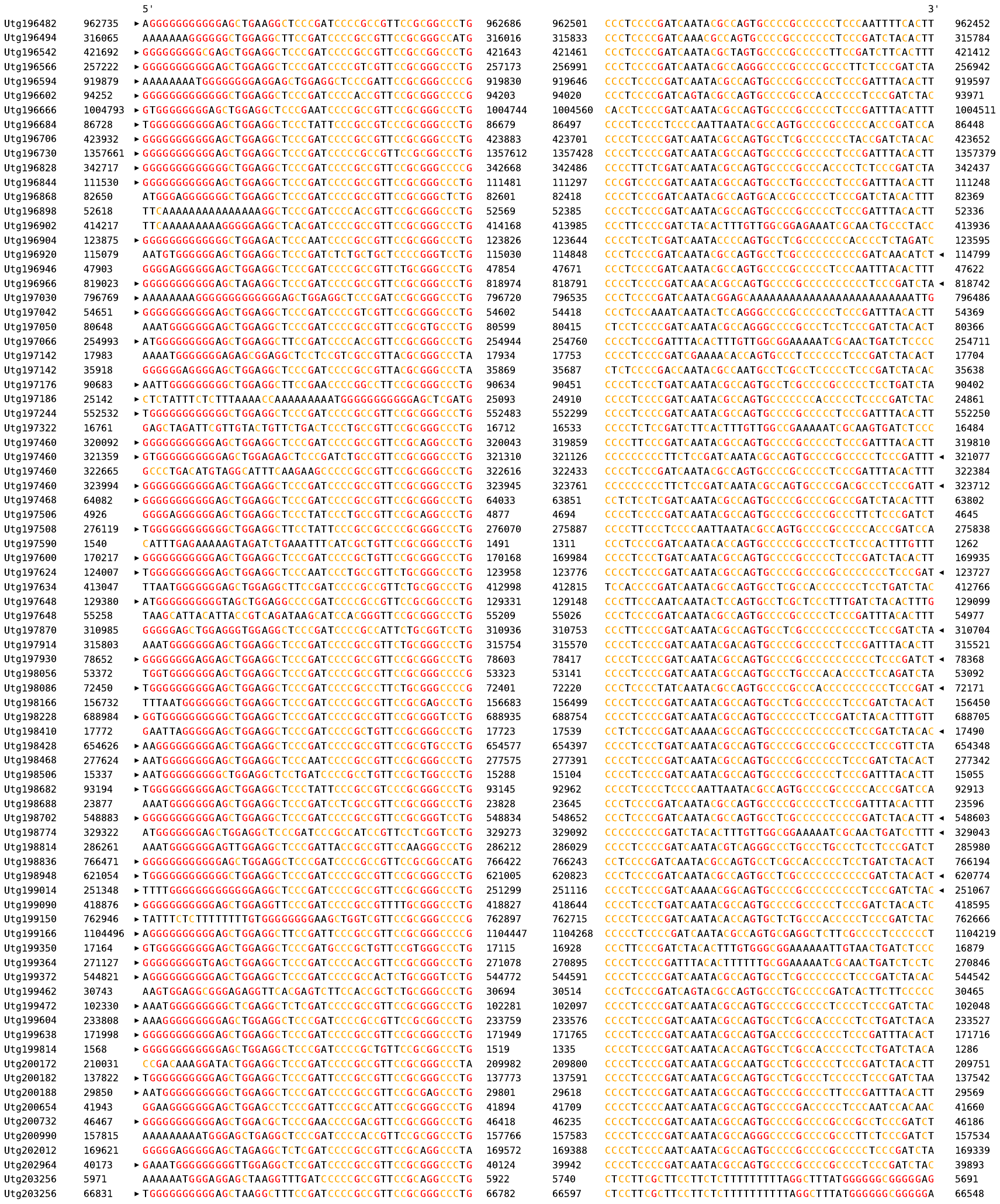
 Fig. S1: Upstream (left) and downstream (right) 50 bp sequences of 185 ArsInsC homologs and their locations. Guanine (G) and cytosine (C) are colored in red and orange, respectively. The region indicated by black arrowhead contains G(C)-stretch.
